# Conserved residues of the immunosuppressive domain of MLV are essential for regulating the fusion-critical SU-TM disulfide bond

**DOI:** 10.1101/2024.06.04.597458

**Authors:** Victoria A. Hogan, Julia Harmon, Miguel Cid-Rosas, Laura R. Hall, Welkin E. Johnson

## Abstract

The ENV protein of murine leukemia virus (MLV) is the prototype of a large clade of retroviral fusogens, collectively known as gamma-type Envs. Gamma-type ENVs are found in retroviruses and related endogenous retroviruses (ERV) representing a broad range of vertebrate hosts. All gamma-type Envs contain a highly conserved stretch of 26-residues in the transmembrane subunit (TM) comprising two motifs, a putative immunosuppressive domain (ISD) and a CX_6_CC motif. The extraordinary conservation of the ISD and its invariant association with the CX_6_CC suggests a fundamental contribution to Env function. To investigate function of the ISD, we characterized several mutants with single amino acid substitutions at conserved positions in the MLV ISD. A majority abolished infectivity, although we did not observe a corresponding loss in intrinsic ability to mediate membrane fusion. Ratios of the surface subunit (SU) to capsid protein (CA) in virions were diminished for a majority of the mutants, while TM/CA ratios were similar to wild type. Specific loss of SU reflected premature isomerization of the labile disulfide bond that links SU and TM prior to fusion and entry. Indeed, all non-infectious mutants displayed significantly lower disulfide stability than wild type MLV Env. These results reveal a role for residues at MLV ISD positions 2, 3, 4, 7, and 10 in regulating a late step in fusion, and suggest that the ISD is part of a larger domain, encompassing both the ISD and CX6CC motifs, that is critical for formation and regulation of the metastable, intersubunit disulfide bond.

**IMPORTANCE:** The gamma-type Env is an extremely prevalent viral fusogen, extensively found within retroviruses and endogenous loci across vertebrate species and are further found in filoviruses such as Ebola virus. The fusion mechanism of gamma-type Envs is unique from other Class I fusogens such as those of IAV and HIV-1. Gamma-type Envs contain a hallmark feature known as the immunosuppressive domain (ISD) that has been the subject of some controversy in the literature surrounding its putative immunosuppressive effects. Despite the distinctive conservation of the ISD, little has been done to investigate the role of this region for the function of this widespread fusogen. Our work demonstrates the importance of the ISD for the function of gamma-type Envs in infection, particularly in regulating the intermediate steps of fusion with the host membrane. Understanding the fusion mechanism of gamma-type Envs has broad implications for understanding entry of extant viruses and aspects of host biology connected to co-opted endogenous gamma-type Envs.

## Introduction

Retroviruses, among other enveloped viruses, use glycoproteins (Env) embedded into the viral membrane to recognize host cell receptors and mediate membrane fusion. Fusion of the viral membrane with the host cell membrane subsequently allows entry of the viral core into the host cell (1). The prototypical class I fusogens are HA of influenza A virus (IAV) and the envelope glycoprotein (Env) of human immunodeficiency virus 1 (HIV-1) (2). However, this overlooks a subset of Class I fusion proteins that use a distinct mechanism for fusion and entry (3–5). These gamma-type Env fusion proteins are found in viruses representing *Alpha-Beta-Delta-* and *Gammaretrovirus* genera. Gammaretroviruses, for which the group is named, have been found to infect fish, amphibians, birds and mammals, indicative of how the boundary between species has been, and likely remains, an easily traversable barrier for these viruses. Gamma-type Envs are further found across host species within endogenous loci (ERVs) (6–8), and these endogenous gamma-type Envs have often been co-opted for a host function (9, 10). An example includes the human Syncytin proteins that play a crucial role in placenta formation (11–14).

The gamma-type Env clade is marked by several distinctive features that are not found in prototypical Class I fusogens such as HIV-1 Env or IAV HA. All gamma-type Envs are trimers composed of three heterodimers each with a surface (SU) and transmembrane (TM) subunit held together by a disulfide bond (3, 15, 16). This specific disulfide bond is unique to gamma-type Envs, and forms between two highly conserved, cysteine-containing motifs (16, 17). Within SU, the CXXC (commonly CWLC) lends its second cysteine to the bond, while the first cysteine is reserved for isomerization during fusion (Figure 1a) (15, 17). Within TM there is a CX_6_CC motif, of which the last cysteine contributes to the intersubunit disulfide bond with SU (17). The other two cysteines in this motif form their own disulfide bond that is thought to be structurally essential for TM. The labile disulfide bond that holds SU and TM together is essential for infectivity and fusion (15). During fusion, after receptor binding, the fusion peptide on the N-terminal side of TM inserts into the host cell membrane. Isomerization of the disulfide bond between SU and TM results om an internal bond within the CXXC motif of SU (15, 18–20). The uncoupling of SU and TM caused by isomerization is believed to allow TM to undergo a final conformational change (referred to as “hair-pinning”) that brings the cellular and viral membranes together to facilitate fusion (1, 21, 22).

**Figure 1:**
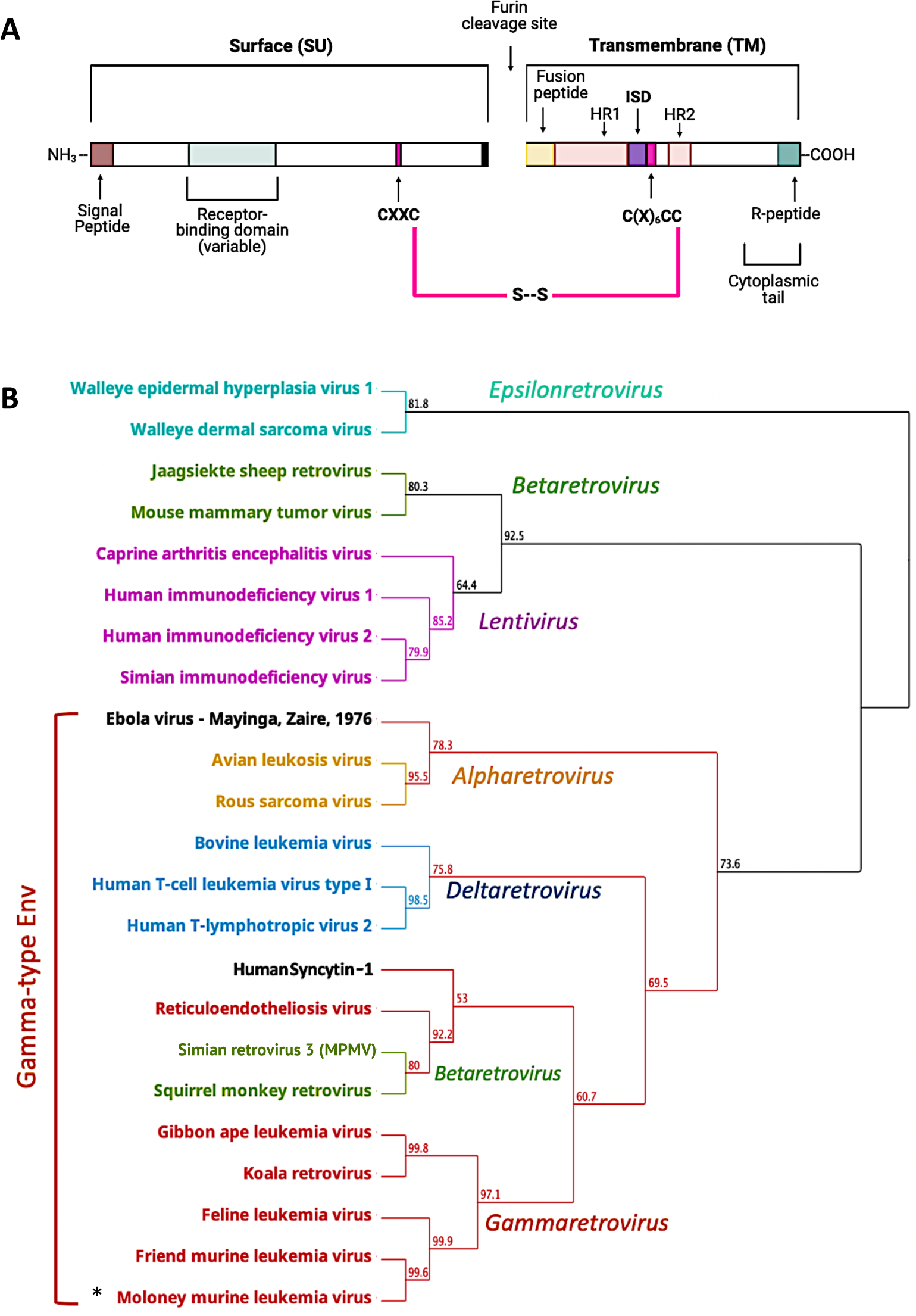
Gamma-type Envs are shared across multiple genera in the retroviridae and beyond, and contain a distinctive disulfide bond between SU and TM. A) Retroviral Env sequence showcasing common features of MLV and related gamma-type Envs. The location of the receptor binding domain is based on MLV however the location of this region can be highly variable and is not completely defined for many Envs. Of note are two cysteine motifs that are joined via a disulfide bond (shown in pink). HR1 and HR2 correspond to the heptad repeats. ISD corresponds to the immunosuppressive domain. B) Geneious Prime (v2022.2.2) generated neighbor-joining tree of envelope proteins using an alignment of TM only, colored by genus except for ebolavirus and syncytin-1. Bootstrap values are shown at branch points. The asterisk corresponds to the gammaretrovirus used for the experiments in this study.

Interestingly, adjacent to the CX_6_CC within TM is another highly conserved region, referred to as the immunosuppressive domain (ISD). Previous work with a small synthetic peptide, homologous to this 17 amino acid stretch just upstream of the CX_6_CC, demonstrated inhibition of lymphocytes and natural killer cells *in vitro* (23–29). Later, work using full Env proteins demonstrated immunosuppressive effects *in vivo* using a murine model of tumor suppression (14, 30–32). This work, combined with the conservation of this region across gamma-type Envs, led to the proposal that the ISD is responsible for a variety of phenomena including viral immune evasion, tumor immune evasion and maternal fetal tolerance (mediated by the syncytins). The literature surrounding the ISD is expansive but still lacks a unifying mechanistic explanation.

Here, we report that the ISD and CX_6_CC motifs are always adjacent, and invariably co-occur with the labile disulfide bond between SU and TM. Additionally, our results suggest that this 26-amino-acid region (spanning both the ISD and CX_6_CC) may have been conserved due to a fundamental role in the regulation and stabilization of the intersubunit disulfide bond between SU and TM, which is crucial in controlling an intermediate step of fusion unique to gamma-type Envs. We specifically demonstrate that small conservative changes within the ISD are sufficient to abolish infectivity, revealing the extreme intolerance for deviation within this region for proper Env function. This is in line with previous literature that demonstrates that larger deletions or double mutations in this region have drastic impacts on infectivity (33, 34). Interestingly, these mutations display little to no impact on the biogenesis of Env and its innate cell-cell fusion capability. Instead, this loss of infectivity is correlated with premature isomerization or instability of the disulfide bond between SU and TM suggesting a role for the ISD in regulating stability of Gamma-type Env proteins.

## RESULTS

### Exceptional conservation of the 26 residues comprising the ISD and CX_6_CC

Transmembrane (TM) subunit based phylogenetic analysis including representatives from all Orthoretroviral genera demonstrate that the Env proteins of alpharetroviruses, gammaretroviruses, deltaretroviruses, and some betaretroviruses constitute a gamma-type Env clade (Figure 1b) (35). The glycoprotein of the filoviruses, including Ebolavirus (EBOV) are also gamma-type entry proteins (Figure 1b). Previously, the ISD was noted to be well conserved in a few frequently studied gamma-type Envs (23). To examine conservation of the ISD in a broader evolutionary context, 36 protein sequences of gamma-type Envs and the EBOV GP, found in NCBI databases, were aligned using Clustal-Omega (Geneious Prime). Envs representing both exogenous viruses from multiple genera and Envs from endogenous retrovirus (ERV) loci were included in the alignment. For the whole alignment the average pairwise identity observed was 17%. In stark contrast to the rest of the Env alignment, a ∼26 residue stretch within TM shows remarkable conservation (Figure 2). This span of conservation holds the residues with the highest positional identity across the whole alignment and contains the only 6 residues with 100% pairwise identity – these correspond to MLV residues N540, R541, D545, and the three cysteines of the CX_6_CC. Several other residues have >75% pairwise identity, including ERV sequences. This region corresponds to the ISD and the CX_6_CC motif, the latter of which is one of two cysteine-containing motifs within Env that form a critical disulfide bond between SU and TM. The length of the ISD and CX_6_CC together also remains constant, indicating that the two motifs were conserved together as part of one 26 residue domain. The labile disulfide bond that forms with the final cysteine of the CX_6_CC is a unique and invariant property of gamma-type Envs that is always seen in conjunction with the presence of the ISD, making both the disulfide bond and ISD two co-occurring properties of gamma-type Envs. It is also worth noting the sequence similarity between Env sequences from both ERVs and actively replicating viruses that represent hosts from many divergent taxa.

**Figure 2:**
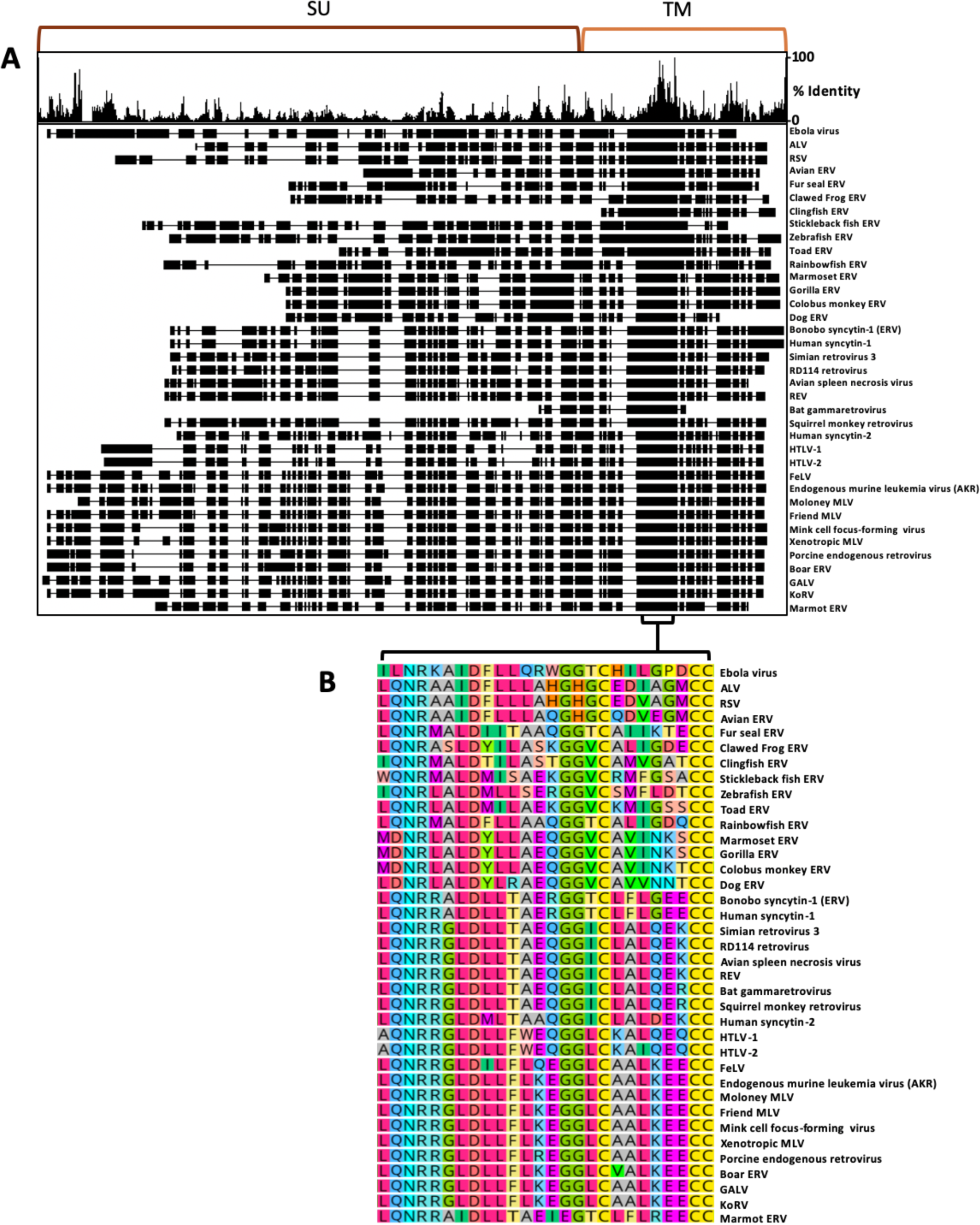
Conservation of the ISD and CX6CC motifs across highly divergent Env sequences. A) Geneious Prime (v2022.2.2) clustal omega amino acid alignment of 37 gamma-type entry protein sequences (where available). Percent identity at each position is shown in the histogram (top). Black boxes indicate aligned residues while horizontal lines indicate gaps in the alignment. Low overall identity is observed for SU and higher for TM. B) The region with the highest conservation and longest unbroken span is shown zoomed in and contains the 26 amino acid stretch that comprises the ISD and CX6CC. ERV = endogenous retrovirus. The diversity of sequences used within the alignment represent hundreds of millions of years of evolution for both viruses and hosts.

Beginning with the leucine previously denoted as the start of the ISD (23), residues L1, Q2, N3, R4, L7, D8, L10, G15 and G16 are remarkably conserved. Most of the remaining positions are less well conserved yet still display strong similarity within phylogenetic clusters. This conservation reveals the crucial importance of these residues, and this region, for the function of the envelope protein which has been maintained over a vast evolutionary time-scale. Previous work demonstrated that certain mutations to the ISD and CX_6_CC (E14R, A20F) reduced infectivity connected with premature interactions with the receptor in the producing cell (14, 32, 34). This, combined with the ISD’s conservation with the disulfide forming CX_6_CC motif, prompted the hypothesis that the ISD may be conserved for an associated function in fusion and disulfide bond stability/regulation.

### Individual mutations at each of the conserved sites of the ISD drastically reduce infectivity

To address the role of the ISD in Env function, several mutants were generated in the murine leukemia virus (MLV) Env using site-directed mutagenesis. Single amino acid changes were made to the most highly conserved residues within the ISD (Figure 3a). All amino acid changes were made in order to minimally impact protein structure, favoring polar to non-polar and charged to polar uncharged when possible. The mutant Q2D was chosen based by replacing the majority residue (glutamine) with one found in a minor subset of Envs (aspartate). All Envs were C-terminally tagged with an Avi epitope tag for western blot analysis.

**Figure 3:**
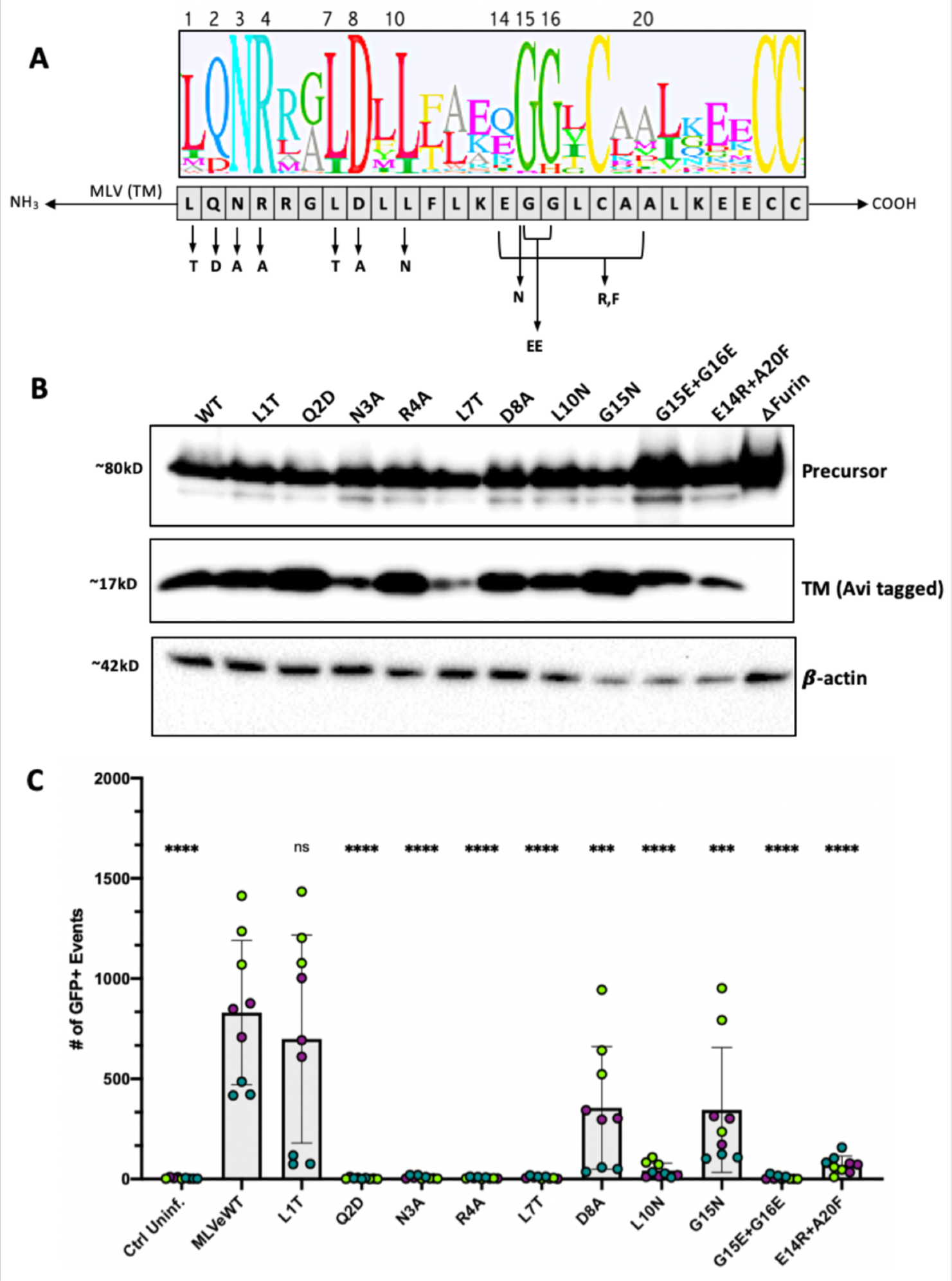
MLV ISD mutants are correctly processed but largely incapable of infection. A) Mutagenic strategy to create 8 single mutants and 2 double mutants to the most conserved regions. *Mutant E14R+A20F is not within the most conserved residues but instead replicates a mutant used previously in the literature (32, 34) believed to have no immunosuppression. Logo plot was based on alignment from figure 2 showing the conserved residues of the ISD. Leucine 1 of the ISD corresponds to L538 in Moloney MLV (accession NC_001501.1) B) Expression via reducing western blot of all Envelope ISD mutants with a furin deletion mutant control to confirm processing phenotypes. All Envs were C-terminally tagged with an Avi epitope tag for visualization. Blots were stained with a rabbit anti-Avi polyclonal antibody. Env processing is demonstrated by the presence of cleaved TM at ∼17kD. A furin deletion mutant was used as a negative control. C) Infectivity of each Envelope mutant pseudotyped with a Mo-MLV core and co-transfected with a packageable GFP plasmid. Virus was produced in 293T cells and murine NIH-3T3 fibroblasts were used as target cells. GFP expression in target 3T3 cells, as measured by flow cytometry, was used as a measure of infection. Three infections, each performed in triplicate, are shown per condition. Data points are colored by infection round (**** = p<0.0001, ns = not significant, Ordinary One-way ANOVA with multiple comparisons).

The glycine at position 15 was eliminated (G15N); however, as this residue is part of a double glycine, it was possible that effects of a mutation here would be masked. Therefore, we additionally included a mutation eliminating both glycines (G15E+G16E). Finally, the double mutant E14R+A20F was also included in our panel. These sites are not highly conserved, however the E14R+A20F mutant was previously reported as abrogating immunosuppression without compromising infectivity (14, 32), while another group reported significant loss in infectivity (34).

All mutant Envs were examined by transient transfection and visualization with western blot of whole cell lysates. A furin cleavage site deletion mutant was used as a negative control for processing. All mutant Envs were expressed and processed to a similar extent as revealed by the presence of a band correspond to the Env precursor polyprotein at ∼80kD and another band corresponding to the cleaved TM subunit at ∼15kD (Figure 3b).

We next tested the ability of the ISD mutants to facilitate infectivity of pseudotyped MLV virions in murine fibroblasts (NIH-3T3 cells). The majority of ISD mutant Envs were incapable of mediating infection. All conditions were lower than wild type MLV Env, except for mutant L1T (L538T in MLV Env), demonstrating that even single conservative changes were enough to drastically impact Env function (Figure 3c). Two mutants showed roughly half the infectivity of wild type, D8A (D545A in MLV Env) and G15N (G552N in MLV Env). As expected, a double glycine mutation had a more pronounced effect on infectivity than G15N alone. In our assays, the previously characterized double mutant, E14R+A20F (32, 34), resulted in significantly reduced infectivity (Figure 3c).

### ISD mutants are still capable of cell-to-cell fusion

We next assessed the intrinsic fusogenicity of each mutant Env in a cell-to-cell fusion assay. In addition, an isomerization deficient mutant was generated by substituting the first cysteine in the CWLC motif (CWLC to AWLC). This eliminates the free thiol of the motif and thus prevents isomerization of the intersubunit disulfide bond (15, 20). Interestingly, all but two of the mutant Envs demonstrated wild type levels of fusion. Only two mutants showed a reduced level of fusion from wild type, Q2D and R4A, although they still showed fusion above control levels (Figure 4).

**Figure 4:**
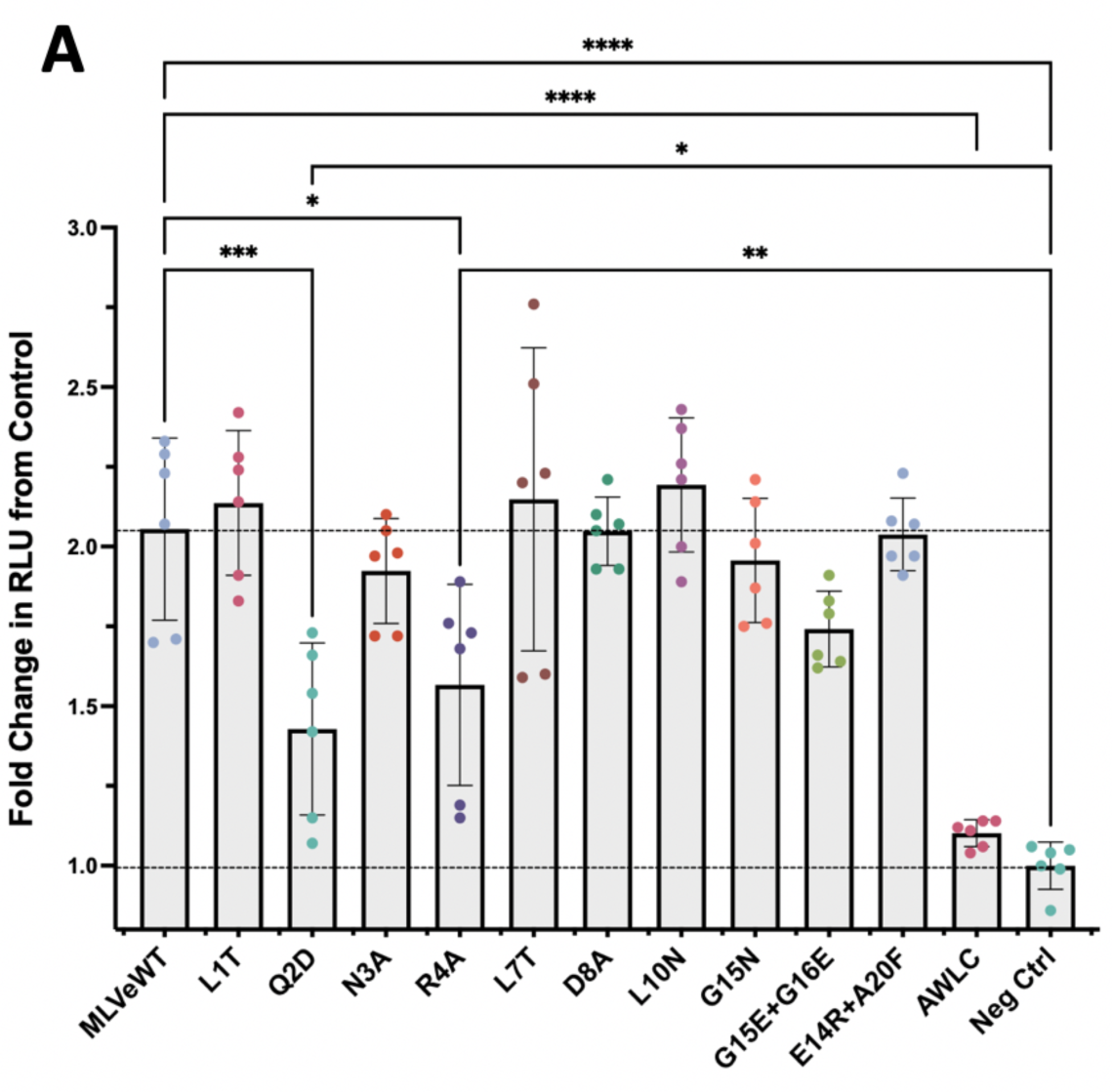
MLV ISD mutants are still fusogenic. A) Cell-to-cell fusion ability of all Env mutants as measured via luciferase assay in which expression of luciferase is turned on when Env mediated fusion occurs (**** = p<0.0001, *** = p<0.001, ** = p<0.01, * = p<0.05, Ordinary One-way ANOVA with multiple comparisons). Lines drawn at wild type mean and negative control mean for visual comparison.

### ISD mutant Envs are incorporated into virions but many have reduced levels of SU

We next investigated the levels of incorporation for each mutant by harvesting viral pellets from the infectious media of producer cells. These viral pellets were separated via SDS PAGE and visualized on a western blot stained with antibodies against SU, a C-terminal tag on TM, and capsid. All Envs were incorporated into virions as shown by the presence of TM (Figure 5a). Compared to wild type MLV Env, N3A and L10N have the lowest ratio of TM to CA, however, the ratio of TM to capsid for all mutants was not significantly different from wild type across multiple experiments (Figure 5b). While we did not observe any major defects in incorporation, certain mutants had lower levels of SU present in viral pellets, particularly N3A, L7T and L10N. In contrast, Q2D consistently displayed wild type levels of SU (Figure 5c).

**Figure 5:**
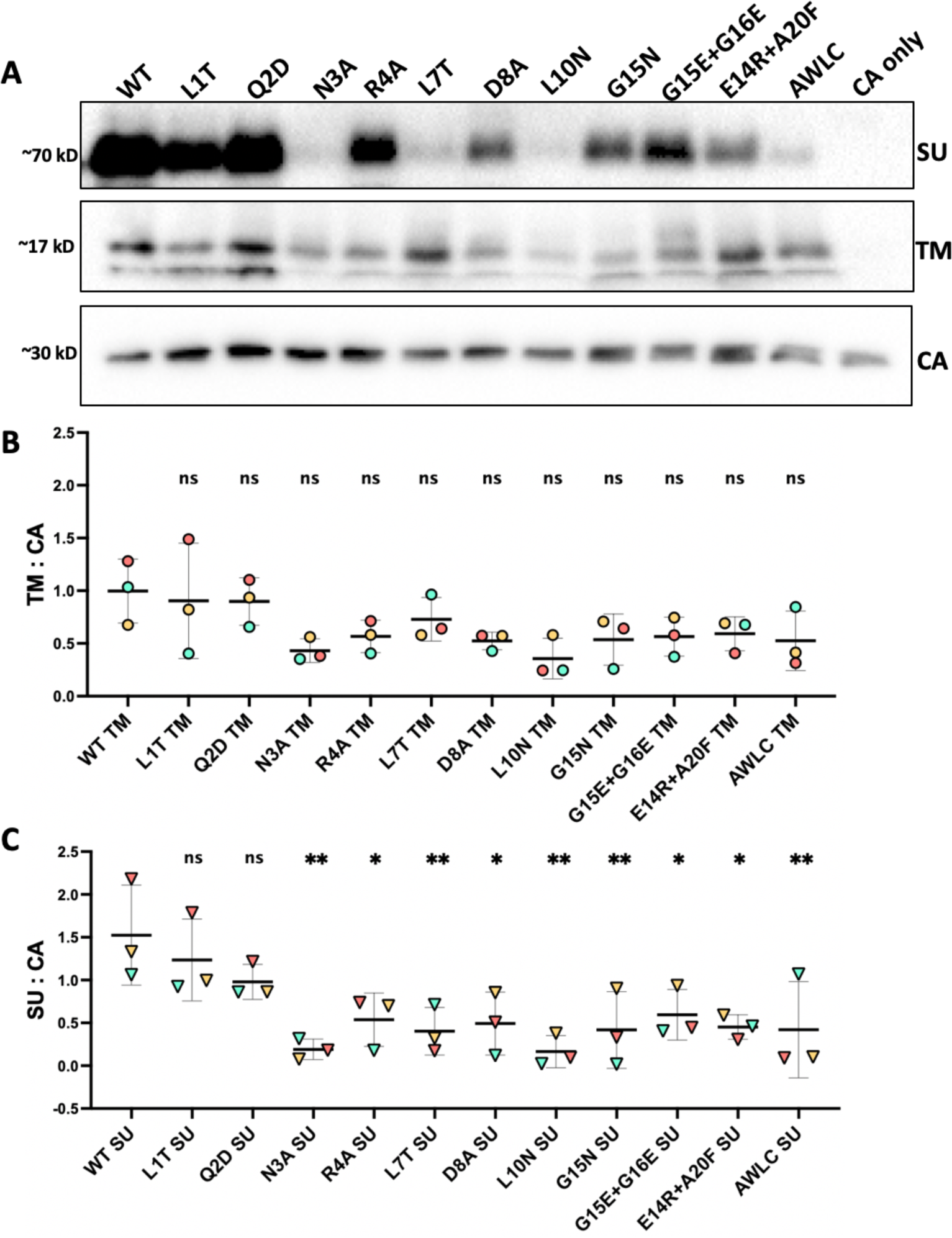
Incorporation of MLV Env ISD mutants into MLV particles. A) Incorporation of Envelope mutants as seen on a western blot of viral pellets. MLV SU was visualized with an anti-MLV-SU antibody, TM was visualized with the same anti-Avi stain used previously, and capsid was imaged using an anti-p30 antibody. B) Ratio of TM to Capsid across three experiments (points colored by experiment). Densitometry was used to quantify levels of CA and TM for three sets of viral pellets. No significant difference was measured between wild type and all mutant conditions (p = 0.09, Ordinary One-way ANOVA with multiple comparisons). C) Ratio of SU to Capsid across three experiments (points colored by experiment). Densitometry was used to quantify levels of CA and SU for three sets of viral pellets. (** = p<0.01, * = p<0.05, ns = not significant. Ordinary One-way ANOVA with multiple comparisons).

The reduced levels of SU suggest that the ISD mutations may destabilize the disulfide bond connecting the two subunits, resulting in loss of SU.

### ISD mutations do not prevent formation of the intersubunit disulfide bond

To confirm the presence of disulfide bond between SU and TM, we ran all mutants on a non-reducing SDS PAGE and used western blot to visualize Env. To confirm that the disulfide bond was forming for each mutant, Env-expressing cell lysates were treated with *N*-Ethylmaleimide (NEM), an alkylating agent which prevents reduction of disulfide bonds. As an additional control, we used the isomerization defective mutant (AWLC) to block dissociation of SU and TM. All samples were harvested in non-reducing conditions to preserve disulfide bonds, and blots were stained with an antibody against a tag on the C-terminus of TM which facilitates visualization of both the Env precursor and the disulfide bonded SU-TM heterodimer (Figure 6).

**Figure 6:**
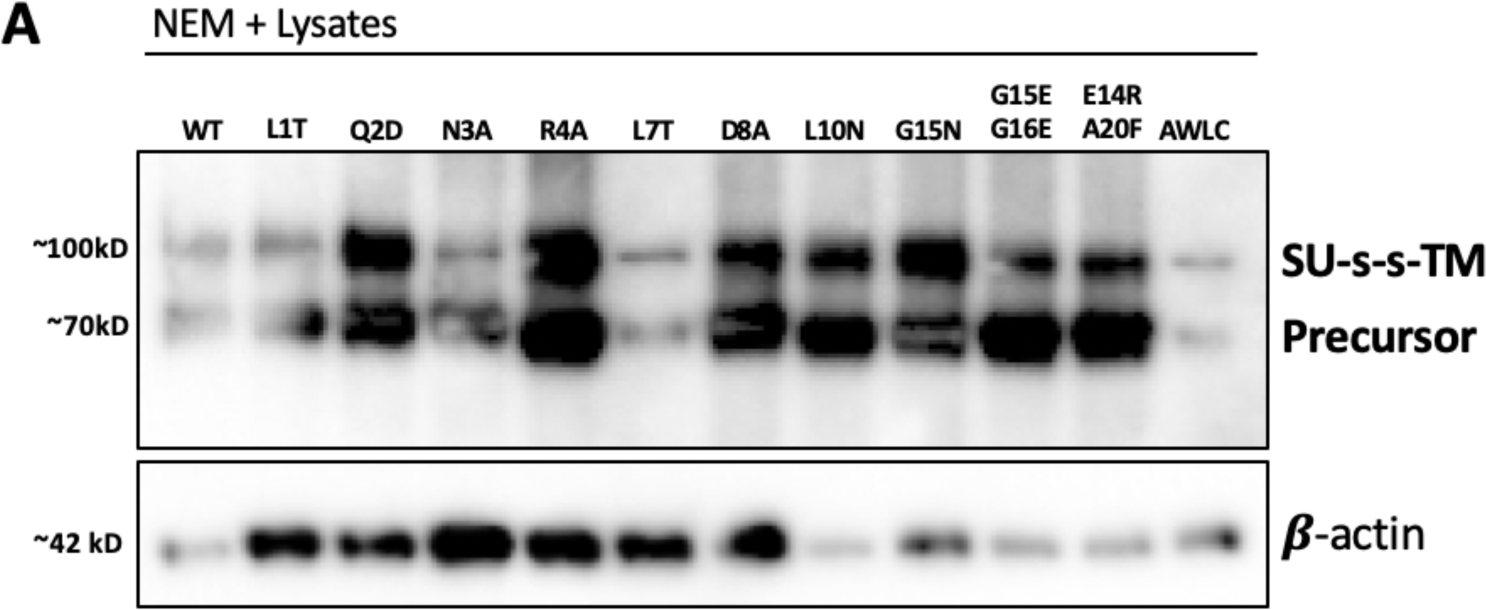
ISD mutations do not prevent formation of the disulfide bond between SU and TM. A) NEM treated whole cell lysates of MLV ISD mutants, stained with anti-Avi to detect TM. Disulfide bonded Env migrates to around 100kD, precursor protein to around 80 kD, SU alone would migrate to 70kD. Beta Actin was used as a loading control for lysates.

A substantial amount of the disulfide intact form was present for wild type. Additionally, for all of the mutants we observed the disulfide bonded form in NEM treated lysates (Figure 6), indicating that the bond was still forming for these mutants.

### Mutations to the ISD promote premature dissociation of SU and TM

To determine the impact of ISD mutations on disulfide stability of Env incorporated into particles, we separated non-reduced viral pellets of all mutants with SDS PAGE and then used western blot to visualize Env with an intact disulfide bond, as well as SU after isomerization of the bond (Figure 7a).

**Figure 7:**
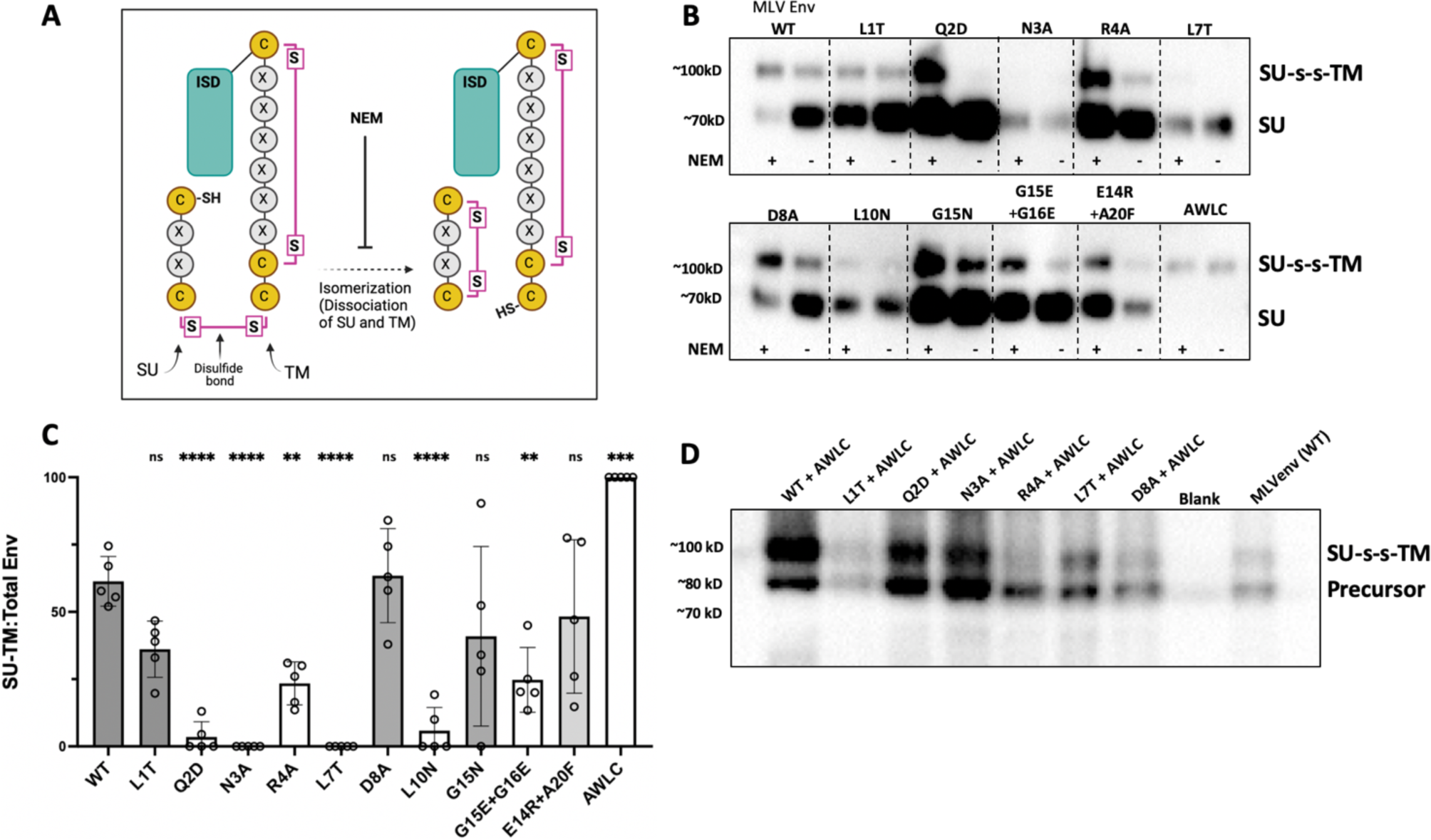
MLV ISD mutants have a less stable disulfide bond. A) Schematic of conserved cysteine containing motifs and formation/isomerization of the disulfide bond between SU and TM subunits. B) Non-reduced viral pellets were separated with SDS PAGE and visualized with western blot. NEM treatment was used to block isomerization of the disulfide bond, essentially fixing any bonded Env present in the samples. Non-treated samples were maintained in non-reducing conditions to preserve intact disulfide bonds. Mutant AWLC no longer has the free cysteine needed to isomerize and acts as a size control for the disulfide bonded form of Env. C) Densitometry of the disulfide bonded band compared to total protein present for untreated supernatant samples across 5 experiments (representative blot shown in panel B). Mutants are colored by infectivity where gray indicates some infectivity above background and white indicates no infectivity above background. D) Western blot of MLV Env ISD mutants combined with an AWLC mutation to block isomerization. Samples were harvested in non-reducing conditions and separated with SDS PAGE.

These blots were stained with an MLV-SU antibody that can distinguish between SU alone, and disulfide bonded SU and TM (Figure 7a). As with the lysates, a substantial amount of the disulfide intact form was visible for wild type, in both NEM and untreated conditions, and no dissociated SU was observed for the AWLC control (Figure 7b).

In contrast, a majority of the mutants, particularly the ones incapable of mediating infection, demonstrated higher levels of SU than disulfide bonded Env. Additionally, these viral pellets allow for quantification of the ratio of disulfide bonded Env to the total Env protein present as a measure of general stability of the disulfide bond. When stability was quantified across 4 independent experiments for the untreated conditions, ISD mutants Q2D, N3A, R4A, L7T and L10N all showed significantly lower levels of the disulfide bonded form in viral pellets when compared to wild type (Figure 7c).

To confirm that the loss of stability seen in our incorporation blots and non-reduced viral pellets was connected to isomerization of the disulfide bond we combined several of the mutants that displayed the least amount of SU in incorporation results with the isomerization resistant mutation AWLC. When combined with this mutation the disulfide bonded form of Env was visible on a western blot (Figure 7d) demonstrating that the mutations to the ISD did not prevent the initial formation of the disulfide bond but that loss of SU seen in Figure 5 was due to early isomerization of the disulfide bond.

## DISCUSSION

The ENV protein of murine leukemia virus (MLV) is the prototype for a large clade of retroviral entry proteins, which we refer to collectively as gamma-type ENVs (6, 36). Gamma-type ENVs are readily identified by the presence of 1) a metastable intersubunit disulfide bond and 2) a highly conserved stretch of 26 residues in the TM subunit. These 26 residues comprise the putative immunosuppressive domain, or ISD (positions 1-17), and a CX_6_CC motif (positions 18-26). The combined ISD+CX_6_CC sequence is found in viruses of the *Alpharetrovirus*, *Gammaretrovirus* and *Deltaretrovirus* genera. It is also found in a subset of viruses in the *Betaretrovirus* genus, including the type-D simian retroviruses (6, 35). Finally, this hallmark sequence is also found in the fusion subunit (GP2) of viruses in the *Filoviridae* family, including Ebola virus (EBOV).

The three cysteines of the CX_6_CC motif are absolutely conserved, reflecting an essential role in formation of two disulfide bonds. One is an intrasubunit bond, which forms between the first and second cysteines, and the other is a metastable, intersubunit bond that forms between the third cysteine and an unpaired cysteine in SU (or GP1, in the case of EBOV). In contrast, the function of the adjacent 17-residue ISD is not well defined. Deletions in the ISD region result in shedding of SU and complete loss of infectivity (33, 37). Consistent with those reports, we found that individual substitutions at highly conserved positions in the MLV ISD result in premature isomerization of the intersubunit disulfide bond. In most cases, the mutation also resulted in reduced levels of virion-associated SU and significant loss of infectivity. Specifically, we observed a correlation between instability of the SU-TM disulfide bond and loss of infectivity (r=0.63, p=0.03, Supp. Figure 1c). Similarly, there was a correlation between shedding of SU and loss of infectivity (r=0.71, p=0.015, Supp. Figure 1d). Based on these observations, we propose that the extraordinary conservation of the ISD reflects a fundamental role in regulating timing of isomerization of the intersubunit disulfide bond during gammaretrovirus fusion and entry.

High-resolution structures of the MLV ENV trimer have not been reported. However, a high-resolution pre-fusion structure of the EBOV GP trimer that includes the ISD sequence has been described (38). Side chains for amino-acids occupying three of the highly conserved ISD residues, at positions 3, 7 and 10, are oriented towards the central axis of the trimer, whereas the side chains of residues at positions 1, 2, 4 and 8 are oriented towards different parts of the associated SU subunit (Figure 8). Alpha fold models suggest MLV Env has a similar TM and ISD structure (Supp. Figure 2). Notably, mutations in the MLV ISD at positions 3, 7 and 10 resulted in nearly identical phenotypes, including destabilization of the disulfide bond, the lowest SU/CA ratios, and loss of infectivity (Table 1). In contrast, the effect of mutations at MLV ISD positions 1, 2, 4 and 8 were variable, perhaps reflecting different interactions with SU. For example, L1T did not detectably destabilize the disulfide bond, did not have a significant effect on infectivity, and had a SU/CA ratio similar to wild type. D8A was similar to L1T, but with a modest reduction in infectivity. Interestingly, Q2D destabilized the disulfide bond while retaining wild type levels of virion-associated SU, and R4A also destabilized the disulfide bond, while retaining intermediate levels of virion-associated SU. Both mutants displayed a reduced capacity for cell-cell fusion as compared to the other mutants. Finally, G15N had only a modest effect on infectivity, while the G15E/G16E double mutant resulted in a complete loss of infectivity. Comparing these observations suggests that different faces of the ISD helix play different roles: for example, based on our results and comparison to the EBOV structure, positions 3, 7 and 10 in the MLV ISD may allow interactions between the TM subunits along the trimeric axis, while some or all of the side chains at positions 1, 2, 4 and 8 may engage in non-covalent interactions between SU and TM.

**Figure 8:**
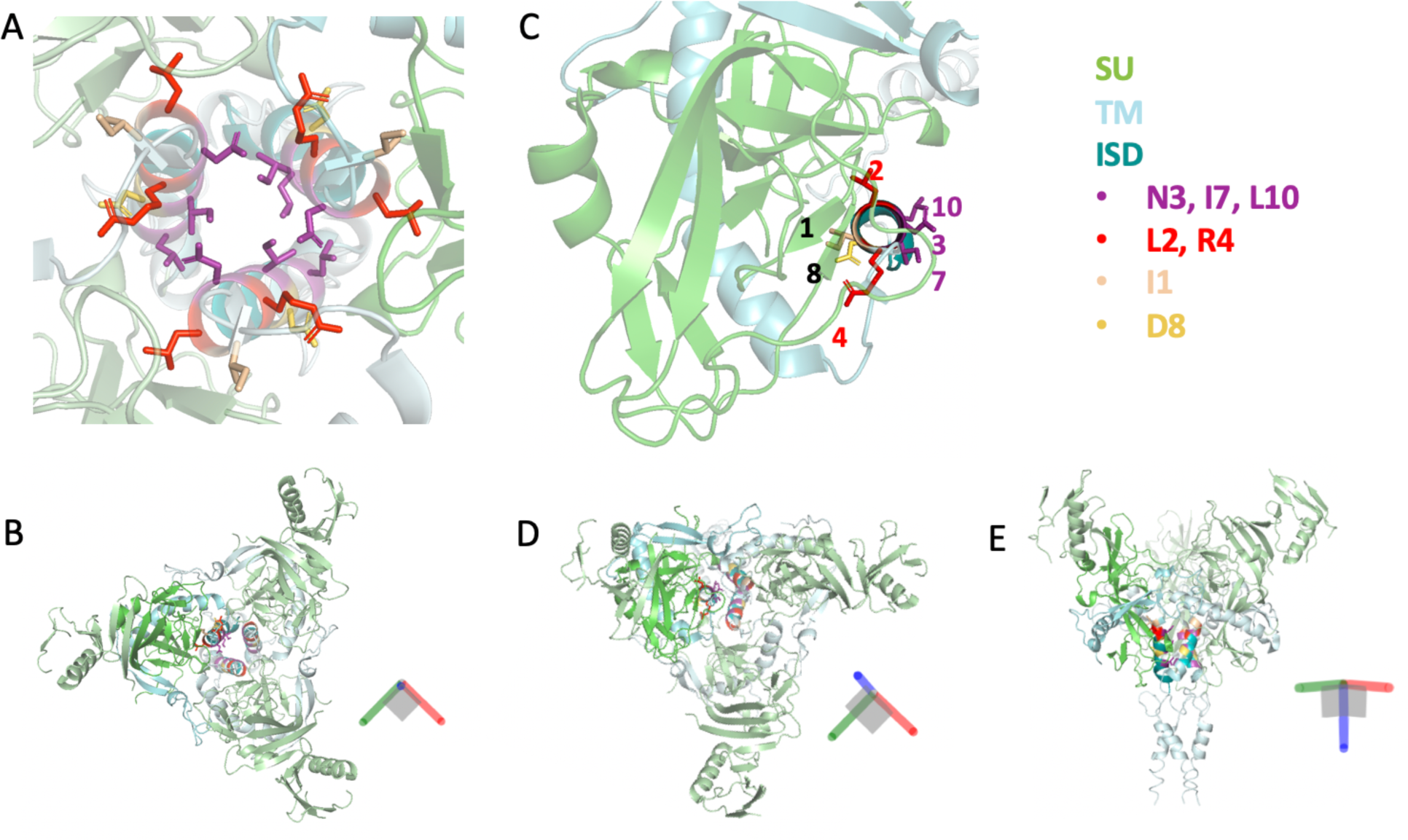
Structure of the pre-fusion Ebola GP trimer highlighting highly conserved residues of the ISD. A) Top down view of the central axis of the pre-fusion structure of Ebola GP (6QD8). Side chains are shown for residues 1, 2, 3, 4, 7, 8 and 10 of the ISD. B) Zoomed out view of panel A. C) Top down view of the alpha helix formed by the ISD. Side chains are shown for residues 1, 2, 3, 4, 7, 8 and 10. For clarity only one heterodimer is shown. D) Zoomed out view of panel C showing the full Ebola GP trimer. E) Side view of the full Ebola GP trimer.

**Table 1:**
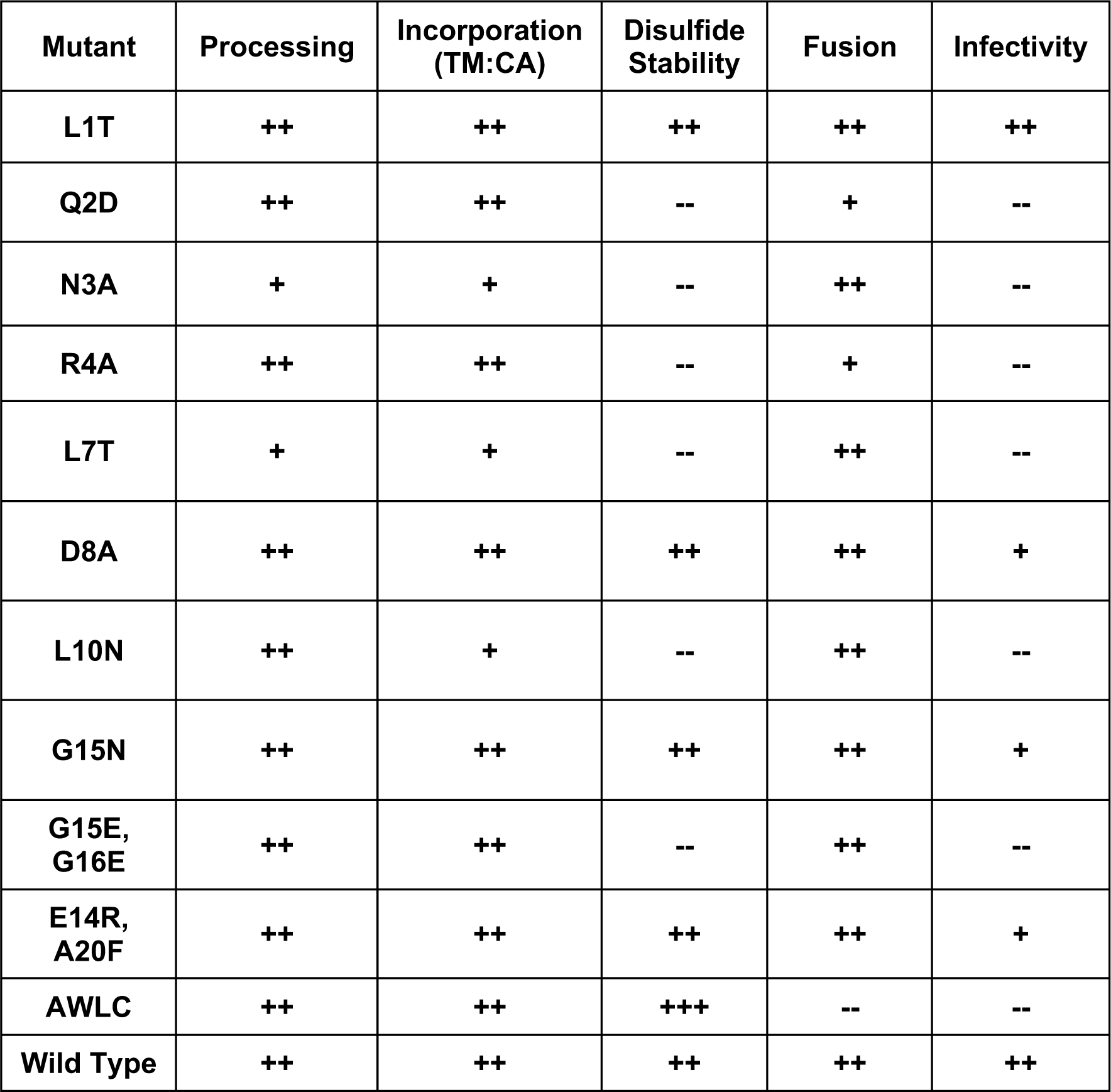
Summary of ISD mutant phenotypes compared to wild type MLV Env. ++ is not significantly different from wild type, – is significantly different from wild type (p<0.05), + is lower than wild type but significantly above negative controls or not significantly different from wild type. AWLC is hyper-stable (unable to isomerize) and therefore listed as +++.

| Mutant | Processing | Incorporation<br>(TM:CA) | Disulfide<br>Stability | Fusion | Infectivity |
| --- | --- | --- | --- | --- | --- |
| L1T | ++ | ++ | ++ | ++ | ++ |
| Q2D | ++ | ++ | -- | + | -- |
| N3A | + | + | -- | ++ | -- |
| R4A | ++ | ++ | -- | + | -- |
| L7T | + | + | -- | ++ | -- |
| D8A | ++ | ++ | ++ | ++ | + |
| L10N | ++ | + | -- | ++ | -- |
| G15N | ++ | ++ | ++ | ++ | + |
| G15E,<br>G16E | ++ | ++ | -- | ++ | -- |
| E14R,<br>A20F | ++ | ++ | ++ | ++ | + |
| AWLC | ++ | ++ | +++ | -- | -- |
| Wild Type | ++ | ++ | ++ | ++ | ++ |

As with influenza A virus HA, the prototypical class 1 fusogen, binding of MLV ENV to its receptor initiates unfolding of TM and insertion of the fusion peptide into the host cell membrane (6). However, in contrast to IAV HA, refolding of MLV TM into the post-fusion structure requires isomerization of the SU-TM disulfide bond, and mutations that “trap” the extended conformation of MLV TM can be rescued by treatment with a reducing agent or by exposure to a soluble receptor binding domain (sRBD) provided *in trans* (22, 39, 40). This is distinct from IAV, where the disulfide linking the HA1 and HA2 subunits remains intact throughout the fusion and entry process (41). Moreover, IAV HA2 does not contain the characteristic ISD and CX_6_CC motifs that define the gamma-type clade of entry proteins. Therefore, the SU-TM bond in gamma-type entry proteins is not functionally related to the IAV HA1-HA2 disulfide bond. In the case of IAV, the bond maintains a stable covalent association between the receptor-binding and fusion subunits, whereas in MLV the bond is metastable and serves as a critical “switch” regulating timing of the final step in fusion and entry (19, 42).

During MLV fusion and entry, reduction of the intersubunit disulfide is coupled to oxidation of the CWLC motif (15, 17, 19). Pinter et al previously noted that the CWLC in MLV SU is analogous to the CXXC active sites of cellular protein disulfide isomerases (PDIs), which also become oxidized during reduction of substrate disulfide (17). Indeed, the intersubunit bond that forms between the CX_6_CC in TM and the CWLC in SU resembles the mixed disulfide intermediate that forms between a PDI active site CXXC and a protein substrate thiol undergoing a coupled reduction/oxidation reaction (17, 43, 44). Our results indicate that the destabilizing effect of mutations in the ISD motif of TM requires the intrinsic isomerization activity provided by the CWLC motif in SU. Therefore, we propose that the ISD specifically functions to premature reduction of the intersubunit disulfide bond by the CWLC. In this model, the cascade of structural rearrangements initiated when SU binds a receptor may serve to relieve ISD-mediated suppression of isomerization, which in turn displaces SU and allows refolding of TM into the post-fusion conformation. Based on extrapolation from the prefusion EBOV trimer structure, we speculate that this involves altering interactions between SU and the ISD helix involving residues 2, 4 and/or 8, and interactions between the three ISD helices involving residues 3, 7 and 10. Comparison of pre- and post-fusion EBOV GP structures suggests that the glycines at positions 15 and 16 in the MLV ISD provide structural flexibility necessary for both the initial extension of TM and, upon isomerization of the bond, its subsequent refolding into the post-fusion, 6-helix bundle.

Alpharetroviruses (e.g. avian leukemia virus, ALV) and filoviruses (e.g. EBOV) represent a variation on the gamma-type ENV (6, 36). These viruses have the labile disulfide bond and the ISD+CX6CC motif, but both lack the CWLC motif found in MLV. One intriguing possibility is that ALV and EBOV depend on cellular PDIs expressed in the endocytosis pathway to reduce the intersubunit disulfide bond. Notably, work on both viruses suggests the existence of an unidentified host factor required for the final step in fusion (19, 45–47). Based on our characterization of the MLV ISD, we propose that the ISDs of ALV TM and EBOV GP2 function to stabilize the intersubunit bonds in these viruses, and that receptor-binding (perhaps in combination with a drop in pH) repositions the ISD such that host-encoded PDIs can access and reduce the disulfide.

As Env is, in a sense, “spring-loaded” for fusion, the control of fusion and protection of the disulfide bond during Env trafficking and virion assembly may be necessary to prevent premature dissociation of the ENV subunits. Multiple mechanisms may exist in Env as controls to ensure fusion occurs only in specific contexts. During viral maturation the MLV protease removes a short peptide from the C-terminal cytoplasmic tail of TM (the R-peptide). R-peptide cleavage results in a significant increase in fusion capability (48–50). At present, it is not clear whether the R-peptide and the ISD regulate different aspects of fusion, or whether they are components of the same regulatory mechanism.

The first reports describing the putative immunosuppressive activity of a synthetic peptide corresponding to the MLV ISD also noted conservation of the sequence in other retroviruses (23). Here, we show that this conservation extends beyond mammalian retroviruses, as evidenced by its presence in endogenous retrovirus (ERV) loci found in the genomes of birds, amphibians and fish. The ISD sequence is also ancient – the sequence is found in human Syncytin-1 and Syncytin-2, which are >25 million years old and 40 million years old, respectively; it is also found in HEMO, another ERV-encoded ENV protein that is >100 million years old, and in the *percomorf* locus of ray-finned fish, which is also >100 million years old (51–54). Given the broad host distribution of ISD-containing viruses among vertebrate species, it is challenging to conceive of an aspect of immunity that would be conserved in birds, fish, amphibians and mammals. The antiquity of the ISD sequence is also at odds with the evolutionary “arms-race” pattern typical of molecular virus-host interactions (55). It is worth noting that the immunosuppressive effects reported thus far have involved a limited number of mammalian retroviruses and mammalian cells (or murine models); therefore, it is possible that the ISDs of these viruses provide an immunosuppressive function in addition to a fundamental role in regulating fusion and entry. However, the latter is a more compelling explanation for the observed conservation of the ISD sequence, and its invariant association with the intersubunit disulfide bond, across hundreds of millions of years of divergent viral evolution. Given the ubiquitous nature of gamma-type Envs and thus the ISD, a deeper understanding of the nature and function of this region and its conservation within Env will be highly valuable to further work on virus-host interactions and the evolution of retroviruses with their hosts.

**Supplemental Figure 1:**
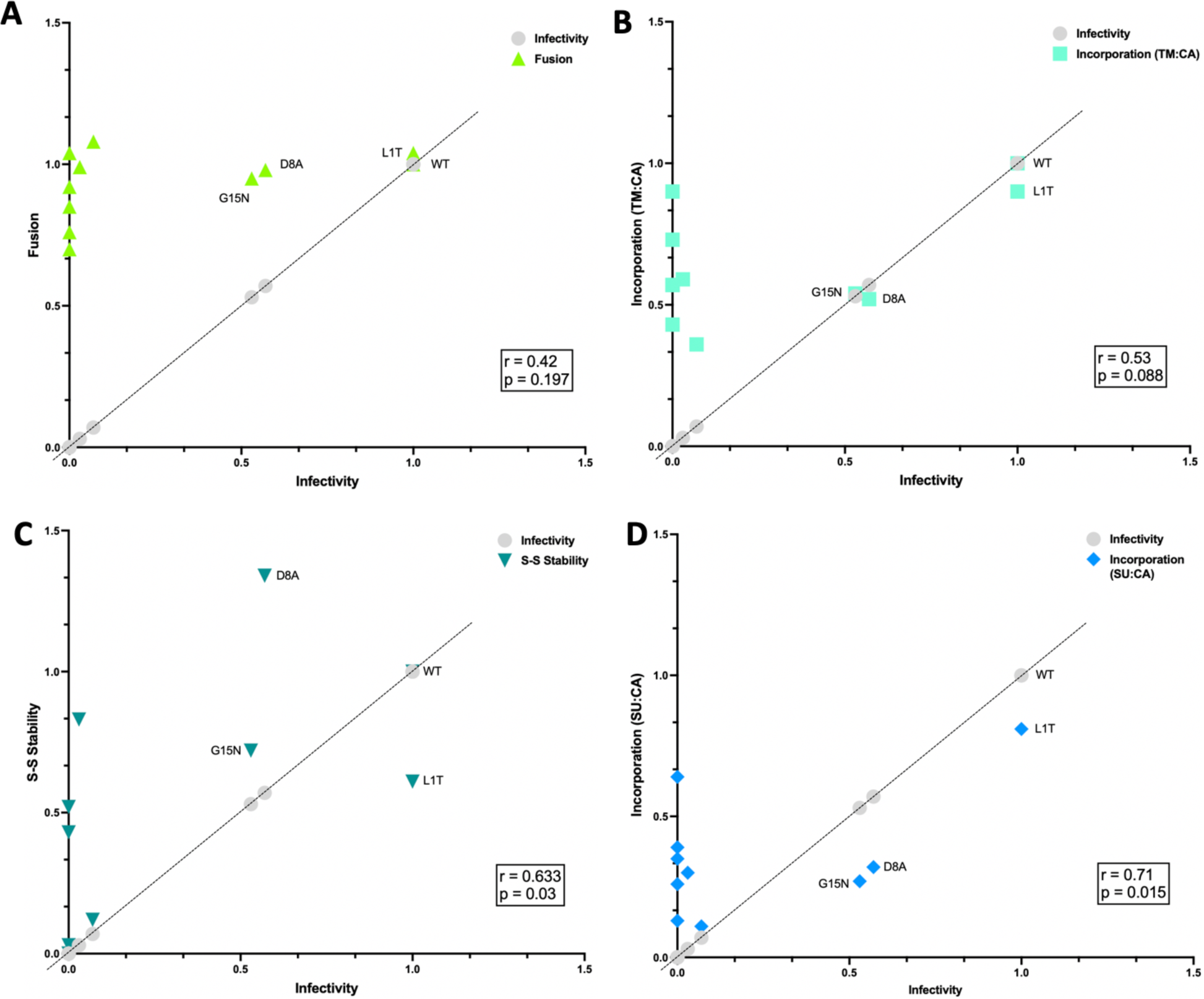
Correlation of ISD mutant phenotypes with infectivity. A) XY data of the correlation between fusion data (Figure 4) and infectivity data (Figure 3c). B) XY data of the correlation between TM:CA data (Figure 5b) and infectivity data (Figure 3c). C) XY data of the correlation between disulfide stability data (Figure 7c) and infectivity data (Figure 3c). D) XY data of the correlation between SU:CA data (Figure 5c) and infectivity data (Figure 3c). All data sets were normalized to wild type.

**Supplemental Figure 2:**
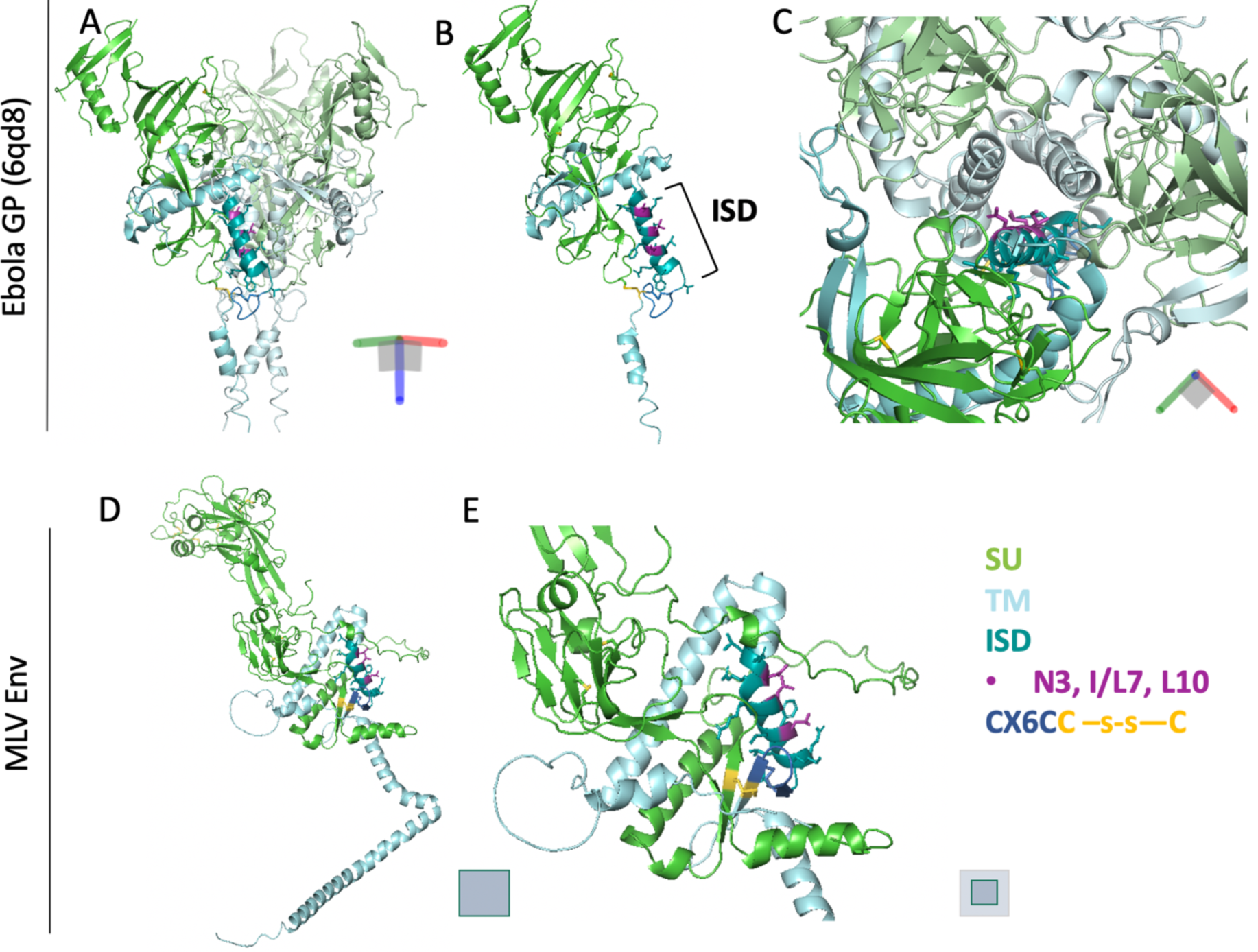
Pre-fusion crystal structures of Ebola GP and Alphafold predictions of MLV Env show conserved structure and position of the ISD. A) A single pre-fusion **heterodimer** of Ebola GP1 and GP2 (6qd8) colored by subunit. Residues N3, I7 and L10 of the ISD are highlighted in purple. B) Pre-fusion trimer of Ebola GP (6qd8) colored by subunit. C) Top-down view of the same structure from panel B, with side-chains visible for the residues of the ISD. Residues N3, I7 and L10 of the ISD are highlighted in purple. D) Alphafold prediction of a “pre-fusion” heterodimer of MLV Env, colored by subunit. E) Zoomed in view of the same structure from panel D to display ISD sidechains. Residues N3, L7 and L10 of the ISD are highlighted in purple.

## METHODS

### Plasmids

Moloney MLV Env was taken from pCL-Eco (Addgene) and placed in a pCGCG backbone upstream of an IRES and GFP via homologous recombination (using New England Biolab’s HiFi Assembly kit) to create ecoMLV-EnvAvi-pCGCG. An Avi epitope tag was appended to the C-terminal end of Env. All subsequent mutants were generated from this plasmid. For infections, pCIG3N (N-tropic MLV gag-pol from Addgene) and pLXIN-GFP were used with Env to produce virus. SIVmac239tat in pBlue and SIVmac239 LTR-Luc in pBlue were used in luciferase fusion assays along with ppMCAT1 (Addgene) to express the MCAT1 receptor.

### Mutagenesis

The Q5 Site Directed Mutagenesis kit from New England Biolabs was used to generate all single and double amino acid mutants. The NEBase changer online tool (https://nebasechanger.neb.com/) was used for mutagenic primer design, and all primers were generated through Integrated DNA Technologies (IDT). All plasmids were grown in TOP10 competent cells and all mutants were verified via forward and reverse sequencing performed by Eton Biosciences.

### Cell Culture

HEK293T cells and NIH 3T3 cells were cultured and maintained using Dulbecco’s Modified Eagle Medium (DMEM) with 10% FBS and 1% Penicillin/Streptomycin and 1% added L-Glutamine. Cells were grown at 37°C. All transfections used Genjet (SignaGen) at a 1:3 ratio of DNA to Genjet reagent and followed the SignaGen Genjet transfection protocol optimized for HEK293T cells.

### Infections and Flow Cytometry

Producer cells were seeded for 60-70% confluence 24 prior to transfection. Transfections followed the SignaGen Genjet transfection protocol. Env (ecoMLV-EnvAvi-pCGCG), MLV gag-pol (pCIG3-N) and a packageable GFP reporter (pLXIN-GFP) were transfected into HEK293T cells at a ratio of 1:2:2 respectively. Transfection mix was left on cells overnight and replaced with fresh D10 18 hours post transfection. After 24 and 48 hours, producer cells were checked for GFP fluorescence to ensure proper transfection. Target cells were seeded at roughly 50,000 cells per well into 12-well plates, 24hrs prior to infection. After 72 hours post-transfection, infectious media was collected from the producer cells and centrifuged for 5 minutes at 1,500 rpm to remove cells and debris. Media was then aspirated off target cells, and 500µL of infectious media was added to each well. This media was then diluted with 500µL of fresh D10 24hrs later. Target cells were harvested for flow cytometry 72 hours post-infection. To harvest cells, each well was trypsinized, resuspended in D10 and pelleted in FACS tubes before being resuspended in 150µL of 4% paraformaldehyde. Each sample was run on a BD FACSAria flow cytometer and subsequently gated for living cells, single cells and GFP expressing cells using FlowJo v8.7.3 BD (Biosciences). An uninfected control was used to establish all gates which were then applied to all samples. Tests for normality and an Ordinary One-Way ANOVA with all conditions compared to wild type were performed using GraphPad Prism software (v.10.0.2).

### Cell-cell Fusion Assay

Receptor null cells were co-transfected with an Env (ecoMLV-EnvAvi-pCGCG) and a SIVmac239 tat plasmid. After 18 hours these cells were resuspended and combined with cells co-transfected with the MCAT1 receptor (ppMCAT1) and an SIVmac239 LTR-luciferase plasmid. Mixed population cell suspension was added in triplicate to a 96-well plate and allowed to grow overnight at 37°C. Cell fusion was measured 24 hours after combining cells via luciferase activity using the Thermo One-Step Glow Assay measured with a Victor plate reader. Raw RLU values were calculated as fold-change from a negative control (SIVmac239 tat transfection only) and an Ordinary One-Way ANOVA was performed with multiple comparisons using GraphPad Prism software (v.10.0.2).

### Western blots

For initial test expressions of Env proteins, HEK293T cells were transfected with Env expressing plasmids using the transfection protocol detailed above. Whole cell lysates were harvested 72 hours post transfection and lysed directly in 2X Laemmli buffer containing 2-Mercaptoethanol. For non-reduced viral pellets, transfection of producers was performed as detailed above. Three days after transfection, media was collected from the producer cells, filtered to remove cellular debris, and centrifuged at 14,000rpm (20,000 x *g*) and 4°C for 3 hours to pellet virions. Viral pellets were then lysed using IP lysis buffer and eluted into 2X Laemmli buffer without 2-Mercaptoethanol or any other reducing agent. All samples were run on 10% polyacrylamide gels at 90V for 2.5 hours in Tris Glycine SDS buffer. Proteins were transferred to a PVDF membrane using 100V for 1.5 hours. Membranes were blocked using 5% milk in 1X PBS and subsequently incubated in primary antibody solution overnight at 4°C. Primary antibody was washed off using 1X PBS with Tween at 0.05% and blots were incubated in HRP-conjugated secondary antibody solution for 1 hour at room temperature. An Enhanced Chemiluminescence kit (Cytiva) was used to image all blots. BioRad Image Lab software (v6.1) was used to perform densitometry for band quantification and all statistical analyses were done using GraphPad Prism software (v.10.0.2) using an Ordinary One-Way ANOVA comparing all conditions to wild type.

### Antibodies

All antibody dilutions were made with 5% milk-PBS. A rabbit IgG anti-Avi polyclonal antibody from Genscript was used at a dilution of 1:3000 in combination with a goat anti-Rabbit IgG-HRP monoclonal antibody at 1:2000. To visualize MLV SU, a goat anti-MLV SU monoclonal (80S-019, Gift from Dr. Monica Roth) was used at 1:2000 in combination with a donkey anti-Goat IgG-HRP secondary antibody at 1:2000. Capsid was visualized with a rat anti-MLV p30 (gag) from Absolute Antibody, and an anti-Rat IgG HRP from Cell Signaling Technologies. A mouse IgG anti-β-actin from Invitrogen was used at 1:3000 in combination with a goat anti-Mouse IgG-HRP conjugated secondary at 1:2000.

### Bioinformatics

All sequences were found on NCBI. Protein alignments and phylogenies were run using Geneious Prime 2022.2.2 (https://www.geneious.com). Accession numbers in order of alignment are as follows:

Ebola virus - Q05320.1, ALV - AFV99542.1, RSV - BAD98245.1, Avian ERV - CAB58112.1, Fur seal ERV - XP_025743503.1, Clawed Frog ERV - QXP50143.1, Clingfish ERV - XP_028321565.1, Stickleback fish ERV - XP_040024727.1, Zebrafish ERV - AF503912, Toad ERV - XP_040289417.1, Rainbowfish ERV - XP_041856078.1, Marmoset ERV - ACI62863.1, Gorilla ERV - AGI61275.1, Colobus monkey ERV - NP_001295060.1, Dog ERV - XP_022270294.1, Bonobo syncytin-1 (ERV) - NP_001291475.1, Human syncytin-1 - Q9UQF0.1, Simian retrovirus 3 (MPMV) - NC_001550.1, RD114 retrovirus - ABS71857.1, Avian spleen necrosis virus - P31796.1, REV - QXV86750.1, Bat gammaretrovirus - AHA85401.1, Squirrel monkey retrovirus - NP_041262.1, Human syncytin-2 - P60508.1, HTLV-1 - Q03816.1, HTLV-2 - AAL30506.1, FeLV - AYG96595.1, Endogenous murine leukemia virus (AKR) - AAB03092.1, Moloney MLV - NC_001501.1, Friend MLV - P26804.1, Mink cell focus- forming virus - AAO37271.2, Xenotropic MLV - AEI59727.1, Porcine endogenous retrovirus - ACD35952.1, Boar ERV - AAQ84, GALV - ALV83307.1, KoRV - ALX81658.1, Marmot ERV XP_015345157.1

Structure predictions were created using Alphafold2 run through ColabFold (https://colab.research.google.com/github/sokrypton/ColabFold/blob/main/AlphaFold2.ip ynb#scrollTo=kOblAo-xetgx), using the MoMLV Env sequence with a 6-residue linker (GAGAGA) between SU and TM after the furin cleavage site. Predicted and crystal structures were graphically analyzed with PyMol (v2.4.2 Schrödinger, Inc.) software. The Ebola GP crystal structure 6QD8 was obtained from RCSB Protein data bank. This crystal structure includes bound antibody fragments which were omitted from our analyses for clarity but may have an impact on the structure of Ebola GP.

## Acknowledgements

We thank Monica Roth for providing the 80S-019 antibodies. We are grateful for the assistance of the Boston College Core facilities, specifically Patrick Autissier Ph.D. who performed the Flow Cytometry used in this work.

This work was supported by discretionary funding provided to WEJ by Boston College. An Eagle Fellowship was provided by the Boston College Career center to MC.

## Author Contributions

VAH designed the experiments. VAH, JH, MC, and LRH performed the laboratory work. VAH conducted data analysis, ran all statistical analyses and prepared all graphical representations. VAH and WEJ wrote the manuscript. All authors critically reviewed the manuscript.

